# ViennaNGS: A toolbox for building efficient next-generation sequencing analysis pipelines

**DOI:** 10.1101/013011

**Authors:** Michael T. Wolfinger, Jörg Fallmann, Florian Eggenhofer, Fabian Amman

## Abstract

Recent achievements in next-generation sequencing (NGS) technologies lead to a high demand for reuseable software components to easily compile customized analysis workflows for big genomics data. We present `ViennaNGS`, an integrated collection of Perl modules focused on building efficient pipelines for NGS data processing. It comes with functionality for extracting and converting features from common NGS file formats, computation and evaluation of read mapping statistics, as well as normalization of RNA abundance. Moreover, `ViennaNGS` provides software components for identification and characterization of splice junctions from RNA-seq data, parsing and condensing sequence motif data, automated construction of Assembly and Track Hubs for the UCSC genome browser, as well as wrapper routines for a set of commonly used NGS command line tools.

## Introduction

Next-generation sequencing (NGS) technologies have influenced both our understanding of genomic landscapes as well as our attitude towards handling big biological data. Emerging functional genomics methods based on high-throughput sequencing allow investigation of highly specialized and complex scientific questions, which continuously poses challenges in the design of analysis strategies. Moreover, the demand for efficient data analysis methods has dramatically increased. While a typical NGS analysis workflow is built on a cascade of routine tasks, individual steps are often specific for a certain assay, e.g. depend on a particular sequencing protocol.

A set of NGS analysis pipelines are available for general [4, 3], and specialized assays such as de-novo motif discovery [6]. While these tools mostly cover the elementary steps of an analysis workflow, they often represent custom-tailored solutions that lack flexibility. Web-based approaches like *Galaxy* [5] cover a wide portfolio of available applications, however they do not offer enough room for power users who are used to the benefits of the command line.

The recently published *HTSeq* framework [2] as well as the *biotoolbox*^1^ package provide library modules for processing high-throughput data. While both packages implement NGS analysis functionality in a coherent manner, we encountered use cases that were not covered by these tools.

## Motivation

The motivation for this contribution emerged in the course of the research consortium “RNA regulation of the transcriptome” (Austrian Science Fund project F43), which brings together more than a dozen experimental groups with various thematic backgrounds. In the line of this project it turned out that complex tasks in NGS analysis could easily be automated, whereas linking individual steps was very labour-intensive. As such, it became apparent that there is a strong need for modular and reusable software components that can efficiently be assembled into different full-fledged NGS analysis pipelines.

We present `ViennaNGS`, a Perl distribution that integrates high-level routines and wrapper functions for common NGS processing tasks. `ViennaNGS` is not an established pipeline per se, it rather provides tools and functionality for the development of NGS pipelines. It comes with a set of utility scripts that serve as reference implementation for most library functions and can readily be applied for specific tasks or integrated as-is into custom pipelines. Moreover, we provide extensive documentation, including a dedicated tutorial that showcases core features of the software and discusses common application scenarios.

Development of the `ViennaNGS` suite was triggered by two driving forces. On the one hand we wanted to return to the open source community our own contribution, which itself is heavily based and dependent on open source software. On the other hand, beside “open science” we advocate for the concept of “reproducible science” [16]. Unfortunately, and to some extent surprising, bioinformatics analyses are often not fully reproducible due to inaccessibility of tools (keyword “in-house script”) or software versions used. It is therefore essential to ensure the entire chain of reproducibility from data generation to interpretation in the analysis of biological data.

## Methods

The major design consideration for the `ViennaNGS` toolbox was to make available modular and reuseable code for NGS processing in a popular scripting language. We therefore implemented thematically related functionality in different Perl modules under the `Bio` namespace (Figure 1), partly building on *BioPerl* [15] and the *Moose*^2^ object framework. Our focus is on consistent versioning, facilitated through Github hosting. In addition, `ViennaNGS` releases are available via the Comprehensive Perl Architecture Network (CPAN), thereby enabling users to get back to previous versions at any time in order to reenact conclusions drawn from shared biological data.

**Figure 1:**
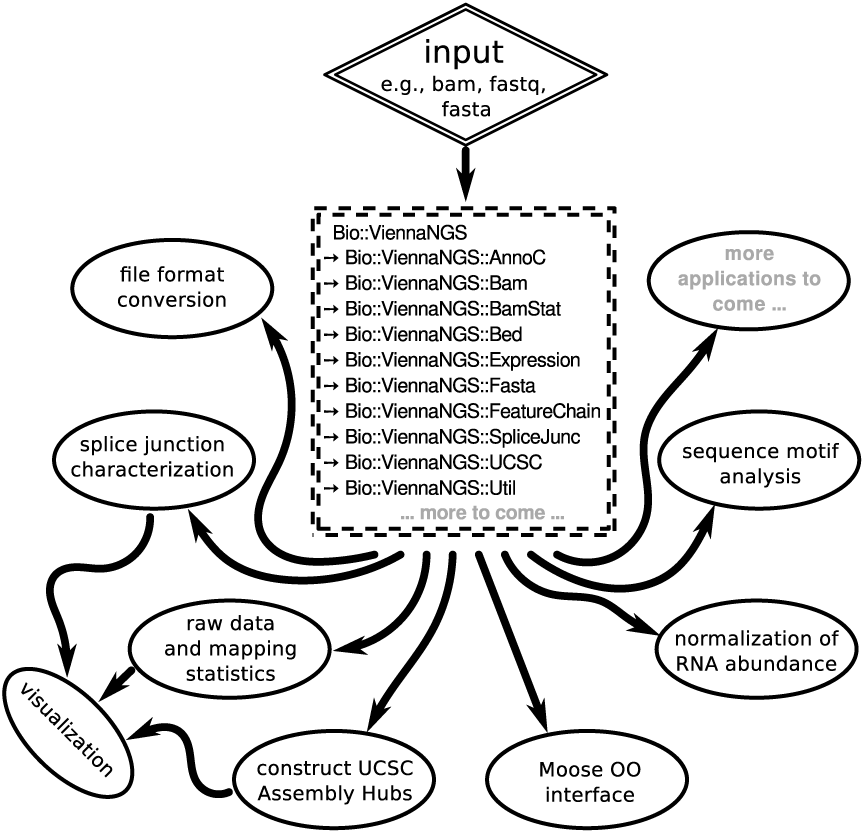
Schematic overview of `ViennaNGS` components. Core modules can be combined in a flexible manner to address individual analysis objectives and experimental setup.

`ViennaNGS` has been designed to close gaps in established analysis workflows by covering a wide range of processing steps from raw data to data visualization. In the following we introduce individual `ViennaNGS` components and describe their main functionality.

## BAM manipulation and filtering

Once mapped to a reference genome, NGS data is typically stored in the widely used SAM/BAM file format. BAM is a binary format, which can easily be converted into text-based SAM format via *samtools* [11] for downstream analysis. However, modern NGS assays produce hundreds of millions of reads per sample, hence SAM files tend to become excessively large and can have a size of several hundred gigabytes. Given that storage resources are always limited, strategies to efficiently retrieve mapping information from BAM format are an asset. To accomodate that, we provide functionality for querying global mapping statistics and extracting specific alignment information from BAM files directly.

`ViennaNGS::BamStat` extracts both qualitative and quantitative information from BAM files, *i.e.* the amount of total alignments, aligned reads, as well as uniquely and multi mapped reads. Numbers are reported individually for single-end reads, paired-end fragments and pairs missing a mate. Quality-wise `ViennaNGS::BamStat` collects data on edit distance in the alignments, fraction of clipped bases, fraction of matched bases, and quality scores for entire alignments. Subsequently, `ViennaNGS::BamStatSummary` compares different samples in BAM format and illustrates results graphically. Summary information is made available in CSV format to facilitate downstream processing.

Efficient filtering of BAM files is among the most common tasks in NGS analysis pipelines. Building on the Bio-SamTools^3^ distribution, `ViennaNGS::Bam` provides a set of convenience routines for rapid manipulation of BAM files, including filters for unique and multiple alignments as well as functionality for splitting BAM files by strand, thereby creating two strand-specific BAM files. Results can optionally be converted to BedGraph or BigWig formats for visualization purposes.

## Genomic annotation

Proper handling of genomic intervals is essential for NGS analysis pipelines. Several feature annotation formats have gained acceptance in the scientific community, including BED, GTF, GFF, etc., each having its particular benefits and drawbacks. While annotation for a certain organism is often only available in a specific format, inter-conversion among these formats can be regarded a routine task, and a pipeline should be capable of processing as many formats as possible.

We address this issue at different levels. On the one hand, we provide `ViennaNGS::AnnoC`, a lightweight annotation converter for non-spliced genomic intervals, which can be regarded a simple yet powerful solution for conversion of bacterial annotation data. On the other hand we have developed an abstract representation of genomic features via generic *Moose*-based classes, which provide functionality for efficient manipulation of BED4, BED6, BED12 and GTF/GFF elements, respectively, and allow for BED format conversion facilitated by `ViennaNGS::Bed`.

`ViennaNGS::MinimalFeature` represents an elementary genomic interval, characterized by chromosome, start, end and strand. `ViennaNGS::Feature` extends `ViennaNGS::MinimalFeature` by two attributes, name and score, thereby creating a representation of a single BED6 element. `ViennaNGS::FeatureChain` pools a set of `ViennaNGS::Feature` objects via an array reference. All intervals of interest can be covered by a `ViennaNGS::FeatureLine` object, which holds a hash of references to `ViennaNGS::FeatureChain` objects (Figure 2).

**Figure 2:**
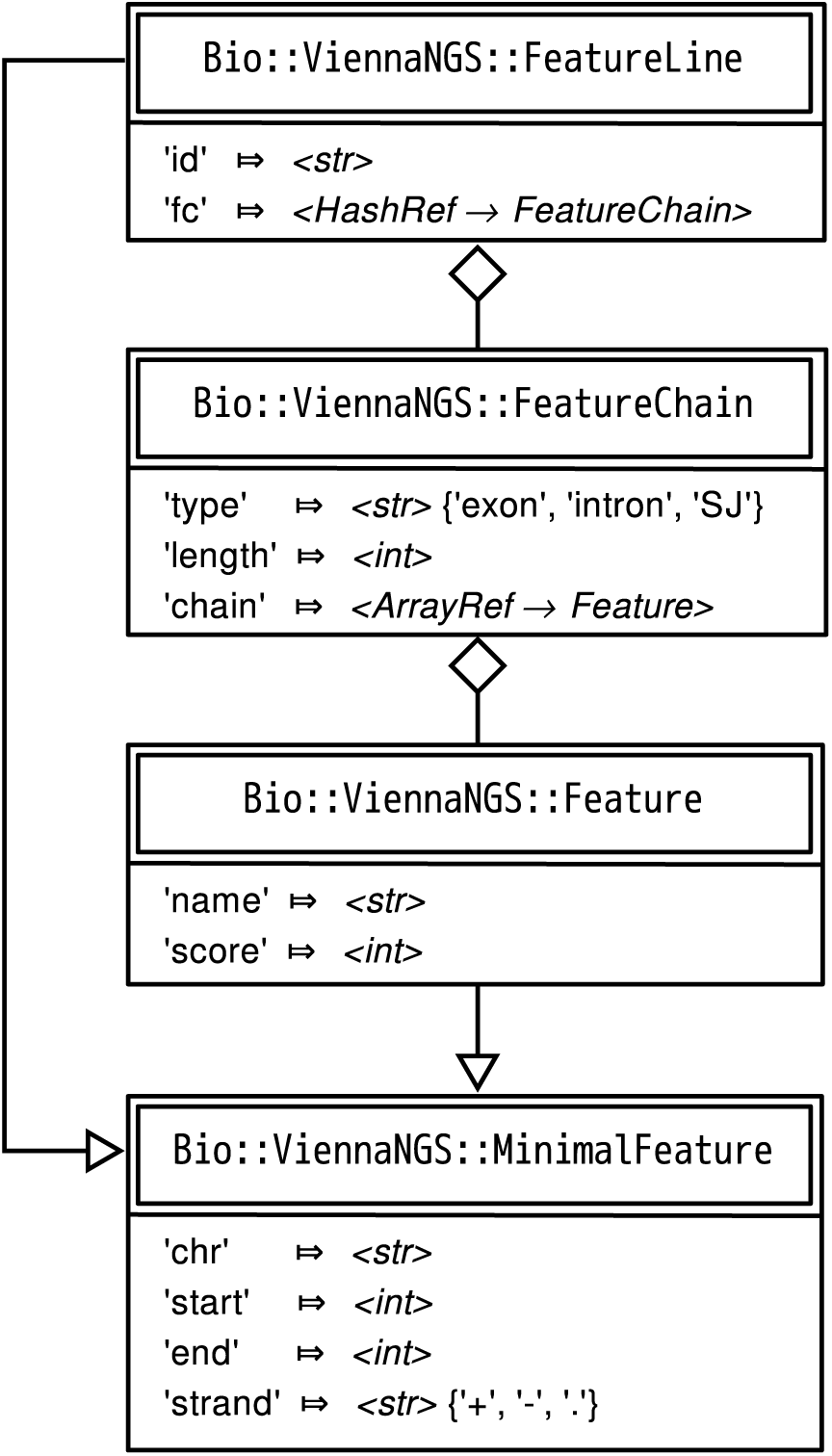
Class diagram illustrating the relations among generic Moose classes which are used as abstract representations of genomic intervals (only attributes are shown).

This framework can handle annotation data by providing abstract data representations of genomic intervals such as exons, introns, splice junctions etc. It allows for efficient description and manipulation of genomic features up to the level of transcripts (Figure 3). Conversely, it is highly generic and can be extended to hierachically higher levels such as genes composed of different transcript isoforms or clusters of paralogous genes.

**Figure 3:**
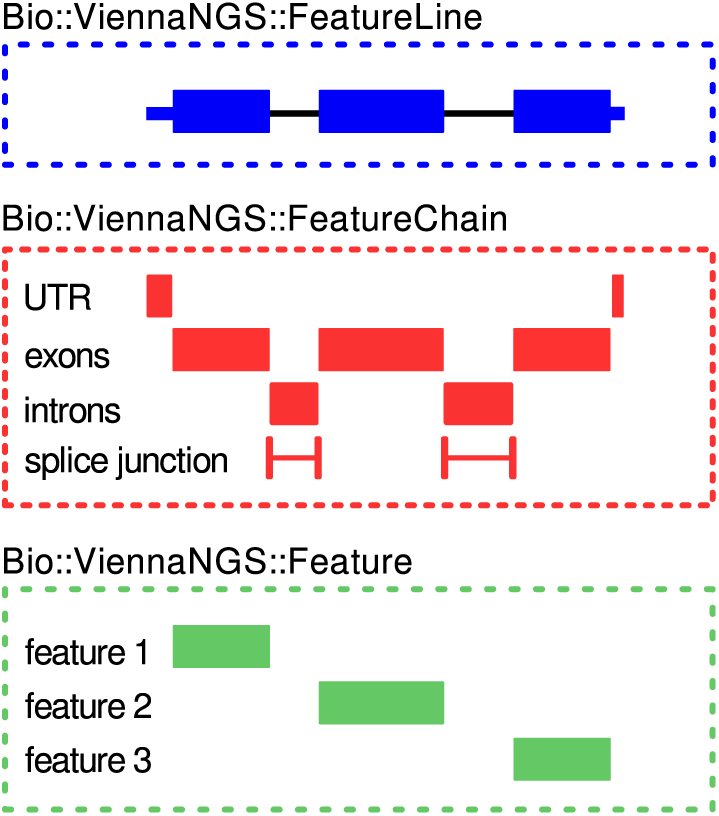
Schematic representation of genomic interval classes in terms of their corresponding feature annotation. Simple invervals (“features”) are charcaterized by `Bio::ViennaNGS::Feature` objects (bottom box). At the next level, `Bio::ViennaNGS::FeatureChain` bundles these, thereby maintaining individual annotation chains for e.g. UTRs, exons, introns, splice junctions, etc. (middle box). The topmost level is given by `Bio::ViennaNGS::FeatureLine` objects, representing individual transcripts.

## Visualization

Another cornerstone of NGS analysis pipelines is graphical representation of mapped sequencing data. In this context a standard application is visualization of Chip-seq peaks or RNA-seq coverage profiles. The latter are typically encoded in Wiggle format, or its indexed binary variant, BigWig, which can readily be displayed within a genome browser. In the same line, genomic annotation and intervals are often made available in BigBed format, an indexed binary version of BED. `ViennaNGS::Util` comes with wrapper routines for automated conversion from common formats like BAM to BigWig or BED to BigBed via third-party utilities [9]. In addition, we have implemented interfaces for a selection of *BEDtools* [13] components as well as a collection of auxiliary routines.

The UCSC genome browser allows to display potentially large genomic data sets, that are hosted at Web-accessible locations by means of Track Hubs [14]. On a more general basis this even works for custom organisms that are not supported by default through the UCSC genome browser, via Assembly Hubs. A typical use case is visualization of genomic annotation, RNA-seq coverage profiles and Chip-seq peaks for *Arabidopsis thaliana* (which is not available through the generic UCSC browser) via a locally hosted Assembly Hub. `ViennaNGS::UCSC` provides all relevant routines for automatic construction of Assembly and Track Hubs from genomic sequence and/or annotation. Besides automated Assembly and Track Hub generation, we support deployment of custom organism databases in local mirrors of the UCSC genome browser.

## Gene expression and normalization

RNA-seq has become a standard approach for gene and transcript quantification by means of measuring the relative amount of RNA present in a certain sample or under a specific condition, thus ideally providing a good estimate for the relative molar concentration of RNA species. Simple “count-based” quantification models assume that the total number of reads mapping to a region can be used as a proxy for RNA abundance [12]. A good measure for transcript abundance is ideally as closely proportional to the relative molar concentration of a RNA species as possible. Various measures have been proposed, one of the most prominent being RPKM (reads per kilobase per million). It accounts for different transcript lengths and sequencing depth by normalizing by the number of reads in a specific sample, divided by 10^6^. It has, however, been shown that RPKM is not appropriate for measuring the relative molar concentration of a RNA species due to normalization by the total number of reads [10, 18].

Alternative measures that overcome this shortcoming have been suggested, e.g. TPM (transcript per million) (eq. 1). Here, rather than normalizing by the total number of mapped reads, a proxy for the total number of transcript samples considering the sequencing reads per gene *r_g_* is used for normalization (eq. 2). The variable rl is the read length and fl*_g_* the feature length of a gene region *g*. Consequently, *T* can be computed by summing over the set of all genes *G*.

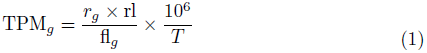

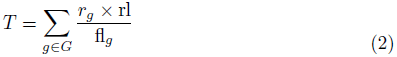

We provide routines for the computation of TPM values for genomic intervals from raw read counts within `ViennaNGS::Expression`.

## Characterization of splice junctions

`ViennaNGS::SpliceJunc` addresses a more specific problem, namely characterization of splice junctions which is becoming increasingly relevant for understanding alternative splicing. This module provides code for identification and characterization of splice junctions from short read mappers. It can detect novel splice junctions in RNA-seq data and generate visualization files. While we have focused on processing the output of *segemehl* [8, 7], the module can easily be extended for other splice-aware split read mappers.

## Documentation and Tutorial

The `ViennaNGS` suite comes with extensive documentation based on Perl’s POD system, thereby providing a single documentation base which is available through different channels, e.g. on the command line via the *perldoc* utility or on the Web via CPAN. Moreover, we provide `ViennaNGS::Tutorial` to guide prospective users through the development of basic NGS analysis pipelines. The tutorial is split into different chapters, each covering a common use case in NGS analysis and describing a possible solution.

## Utilities

The `ViennaNGS` suite comes with a collection of complementary executable Perl scripts for accomplishing routine tasks often required in NGS data processing. These command line utilities serve as reference implementations of the routines implemented in the library and can readily be used for atomic tasks in NGS data processing. Table 1 lists the utilities and gives a short description of their functionality.

**Table 1:** Overview of the complementary utilities shipped with `ViennaNGS`. While some of these scripts are re-implementations of functionality available elsewhere, they have been developed primarily as reference implementation of the library routines to help prospective `ViennaNGS` users getting started quickly with the development of custom pipelines.

| Utility | Description |
| --- | --- |
| `assembly_hub_constructor.pl` | Construct Assembly Hubs for UCSC genome browser visualization |
| `bam_quality_stat.pl` | Compute mapping/quality statistics along with publication-ready figures |
| `bam_split.pl` | Split BAM files by strand |
| `bam_to_bigwig.pl` | Produce BigWig coverage profiles from BAM files for visualization |
| `bam_uniq.pl` | Filter uniquely and multi mapped reads from BAM files |
| `bed2bedGraph.pl` | Convert BED to (strand specific) bed-Graph format |
| `extend_bed.pl` | Extend genomic intervals in BED format at the 5', 3', or both ends |
| `gff2bed.pl` | Convert bacterial RefSeq GFF3 annotation to BED12 format |
| `kmer_analysis.pl` | Count k-mers of predefined length in FastQ and Fasta files |
| `MEME_xml_motif_extractor.pl` | Compute basic statistics from MEME XML output |
| `newUCSCdb.pl` | Create a new genome database in a local UCSC genome browser instance |
| `normalize_multicov.pl` | Compute normalized expression data in TPM from read counts |
| `sj_visualizer.pl` | Convert splice junctions in segemehl BED6 splice junction format to BED12 |
| `splice_site_summary.pl` | Identify and characterize splice junctions from RNA-seq data |
| `track_hub_constructor.pl` | Construct Track Hubs for UCSC genome browser visualization |
| `trim_fastq.pl` | Trim sequence and quality fields in FastQ format |

## Discussion

`ViennaNGS` is a comprehensive software library for rapid development of custom NGS analysis pipelines. We have successfully applied its components in the course of an ongoing, large scale collaboration project focusing on RNA regulation. It has been used with different genomics assays in a wide range of biological systems, including human, plants and bacteria. While we have primarily applied `ViennaNGS` in combination with the short read aligner *segemehl* [8, 7], it has also been used with *Tophat* [17] output very recently in a large scale transcriptome study of Ebola and Marburg virus infection in human and bat cells (Hölzer et al., unpublished data). Moreover, `ViennaNGS` will be used for automated UCSC genome browser integration in an upcoming version of TSSAR [1], a recently published approach for characterization of transcription start sites from dRNA-seq data.

`ViennaNGS` is actively developed and its functionality is constantly extended. In this line, we encourage the scientific community to contribute patches and novel features.

## Data availability

Input data for the `ViennaNGS` tutorial is available from http://rna.tbi.univie.ac.at/ViennaNGS

## Software availability

The `ViennaNGS` distribution is available through the Comprehensive Perl Architecture Network (CPAN) at and GitHub.

1. http://search.cpan.org/dist/Bio-ViennaNGS
2. https://github.com/mtw/Bio-ViennaNGS
3. Software license: The Perl 5 License

## Author contributions

MTW, JF, FE, FA designed and implemented the software. MTW and FA wrote the manuscript. All authors approved the final manuscript.

## Competing interests

No competing interests were disclosed.

## Grant information

This work was funded by the Austrian Science Fund (FWF projects F43 to MTW, FA and FE) and the Research Platform “Decoding mRNA decay in inflammation” by the University of Vienna to JF.

## Footnotes

1 https://code.google.com/p/biotoolbox

2 https://metacpan.org/pod/Moose

3 https://metacpan.org/release/Bio-SamTools

